# Estimation of time to contact in lateral motion and approach motion

**DOI:** 10.1101/408310

**Authors:** Asieh Daneshi

## Abstract

The ability to estimate precisely the time to contact (TTC) of the objects is necessary for planning actions in dynamic environments. However, this ability is not the same for all kinds of movement. Sometimes tracking an object and estimating its TTC is easy and accurate and sometimes it is not. In this study, we asked human subjects to estimate TTC of an object in lateral motion and approach motion. The object became invisible shortly after movement initiation. The results proved that TTC estimation for lateral motion is more accurate than for approach motion. We used mathematical analysis to show why humans are better in estimating TTC for lateral motion than for approach motion.

## 1 Introduction

In everyday life, there are many situations that require us to either avoid or intercept a moving object, even when they are not continuously in view. These objects may approach the observers or just pass laterally, from one side to the other, in front of them. Examples of approach motion include hitting or catching a ball or driving in a street alongside other vehicles, while confronting vehicles when crossing a street is an example of lateral motion (we assume that the individual is headed perpendicular to the direction of moving cars). Estimation of time to contact (TTC), which is the time it takes for an object to reach an observer or a particular place, is critical in these situations.

During the past decades, several studies have been performed to understand different aspects of how humans and animals estimate time to contact. However, there are still a lot of unanswered questions. One of these questions is what causes the difference between TTC estimation for approach motion or lateral motion. In this study, first we conducted a simple experiment in a 3D environment to examine different aspects of TTC estimation in approach motion and lateral motion. Then, we used mathematical analysis to explain the results.

The ratio of an object’s size to its rate of expansion, called tau, is proposed as a prominent source of information for TTC judgments [1, 2].

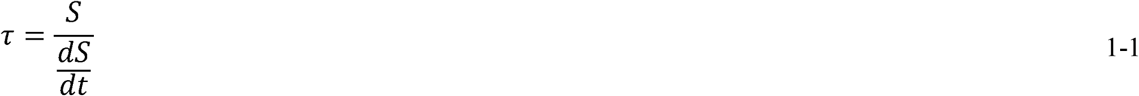

While this provides a reliable judgement of TTC in many situations, it has been shown that a number of additional sources of information are used by the observers, such as binocular disparity [3-5] and vertical velocity [6].

Later, an extension of tau, termed ‘tau-margin’ was formulated to encompass changes in both angular size (looming), *θ*, and the angular gap size, *φ* (see Figure 1). According to the formula that they presented, time-to-arrival can be specified as (Bootsma & Oudejans, 1993):

**Figure 1.**
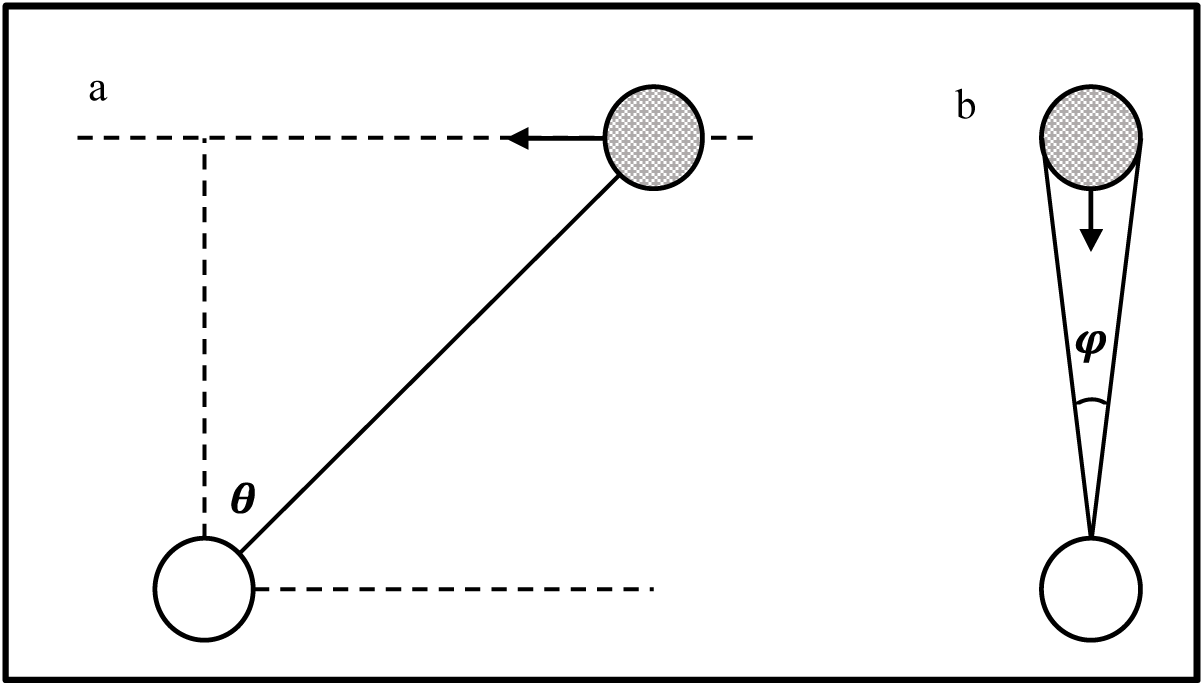
Optical variables computed and used by the model for estimating time-to-contact.

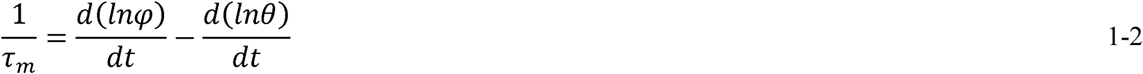

This is a general equation that simplifies to the TTC condition as proposed by [1] when the object moves on a head-on trajectory 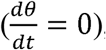, and to the simple 2D gap closure condition when the object does not expand 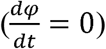. It has been shown that observers are sensitive to the combination of these optical variables (Phi and theta), though with unequal weighting [7]. Later, it was found that observers are sensitive to both the expansion and orientation components of object motion trajectories, including observer self-motion [8].

The formulation of tau is based on a first-order description of object velocity and thus does not consider accelerations. It has been shown that observers perform interceptive actions based on only the linear estimate of tau, even when confronting accelerating objects [9]. It is shown that in general, observers are poor at accounting for accelerations, supporting the idea that judgments of gap closure (see Figure 1-a) are also based on a first-order tau estimate [10]. A similar result was reported in (TTC) estimation, in which the object moved in depth but not on a collision path to the observer [11, 12]. Altogether, these results propose that in general, time-to-contact estimations are based on combination of unambiguous first-order angular velocity estimates. However, there are a few studies addressing the implications or use of a combined-cue tau computation for estimating TTC for all motion conditions.

In 2005, a functional imaging study was conducted to find out the brain areas activated during estimating TTC of approach motion and lateral motion (called looming and gap closure tasks in that study). The researchers suggested the presence of separate mechanisms for these tasks [13]. The tasks that are often considered in laboratory TTC studies can be classified into two main groups: coincidence anticipation (CA) and relative judgment (RJ) tasks. In CA tasks, participants should make a simple response (e.g. press a button) when the moving object reaches a particular place, called contact point [14]. In an important type of CA tasks, often referred to as prediction motion (PM) tasks, the moving object disappears before reaching the contact point or hides behind a cover. Then, participant must do a simple action (e.g. press a button) temporally coincident with the time that is expected for the moving object to reach a specific point. The PM paradigm is used as a relatively straight method to assess individuals’ ability to estimate absolute TTC (e.g. [14]). The main purpose of PM tasks is to understand which visual information observers use to estimate TTC. To this aim, task design variables related to object’s motion (e.g., velocity, extrapolation distance and/or duration) could be manipulated. Most PM studies have focused on lateral motion. One of the first studies which has considered nonlateral motion[15], found that the accuracy of TTC judgments in a PM task decreases as actual TTC increases, and errors typically consist of underestimations. Furthermore, TTC judgments were more accurate with lateral motion than with approach motion D [15]. Later, other researchers obtained similar results when they were studying the contribution of perception and cognition in TTC judgment [16, 17]. They also found that accuracy of TTC estimation is higher for lateral motion than for approach motion [18]. In 1993, researchers evaluated the parameters that influence contact-time judgments and concluded that the position and speed are important in TTC estimation but they are not the only parameters that individuals use to estimate TTC. The tau parameter which is the ratio of visual angle to its derivative through the time 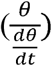 better explains the TTC estimation. They also found that the decision process in their task is dominated by distance information rather than the speed of the target. Recently, several PM studies approved that an increase in actual TTC decreases the accuracy of TTC judgment [19-21]. However, their main goal of these studies was to study the role of attention in estimating time to contact of one or more objects, and have focused only on lateral motion. In 2015, an experiment was conducted to evaluate contribution of velocity and distance to time estimation during self-initiated time-to-contact judgment [21]. In the task that they designed, a ball moved horizontally and the participants were asked to launch a stationary ball in a vertical direction towards the first ball, so that they collide. In this study participants had to estimate a kind of nonlateral motion. However, the experiment was 2-dimensional (2D) and did not represent approach motion. Indeed, the participants viewed the experimental space from the top view (2D); therefore, however the motion trajectories of the balls were perpendicular to each other, this could not be categorized as an approach motion. But, in our study we simulated a virtual 3-dimensional environment so that the participants could perceive lateral motion and approach motion well.

## 2 Materials and methods

### 2.1 Experimental apparatus

#### a. Subjects

Twenty students from Amirkabir University of Technology-Tehran Polytechnic (10 women, 10 men, age 24.85 years±3.80 (mean±SD), min age 19, max age 31) participated in these tests voluntarily. All participants had normal or corrected-to-normal visual acuity. They were healthy and without any known oculomotor abnormalities. Participants were not informed of experimental hypothesis and gave written informed consent to their participation in the experiment.

#### b. Experiment condition

Participants sat on a chair facing a 17″ computer display located at a viewing distance of approximately 50 cm, in a room with normal light. Stimuli were generated with Unity3d, and presented on a desktop computer equipped with a 2.90 GHz Intel Corei7 processor. The screen resolution was 1920×1080 pixels (horizontal by vertical) and the display rate was 60 Hz.

### 2.2 Experiment 1

In the first section (hereafter referred to as “lateral paradigm”), time-to-contact (TTC) estimates for a target car (3 cm length, 1.3 cm width, 1.1 cm height) moving at constant velocity in frontoparallel plane from right to left were obtained using a prediction motion (PM) task (see [19]). Figure 2 shows a screenshot of one of the trials in this experiment. The constant velocities were randomly selected from three values: 3 cm/s, 4.5 cm/s, 6 cm/s. After 2 seconds, the car became invisible. The point that the car disappeared was the same in all trials, so the initial position of the car was arranged so that the visible time was 2 seconds for all trials (for 3 cm/s, 4.5 cm/s, 6 cm/s the distance between initial position of the target car and disappearance point was respectively 6 cm, 9 cm, 12 cm). The car did not reappear after it became invisible. To make it possible to study the influence of distance between the observer and the contact point on TTC estimation, observation point was randomly set at a distance of 0 cm, 3 cm, or 6 cm from the red line. Participants were asked to press a response key (down arrow key) at the instant when the car would have reached a red line placed in its path at a 9 cm distance from the invisibility point. No feedback on TTC estimation error was provided. There was a two-second time interval between two consecutive trials. **Error! Reference source not found.** shows a schematic of the lateral paradigm. Table 1 shows the values of the parameters in the figure. Nine combinations were generated from three velocity values for the target car and three distance values between the observation point and the red line. Each trial was presented 10 times in random order, for a total of 90 trials.

**Table 1.**
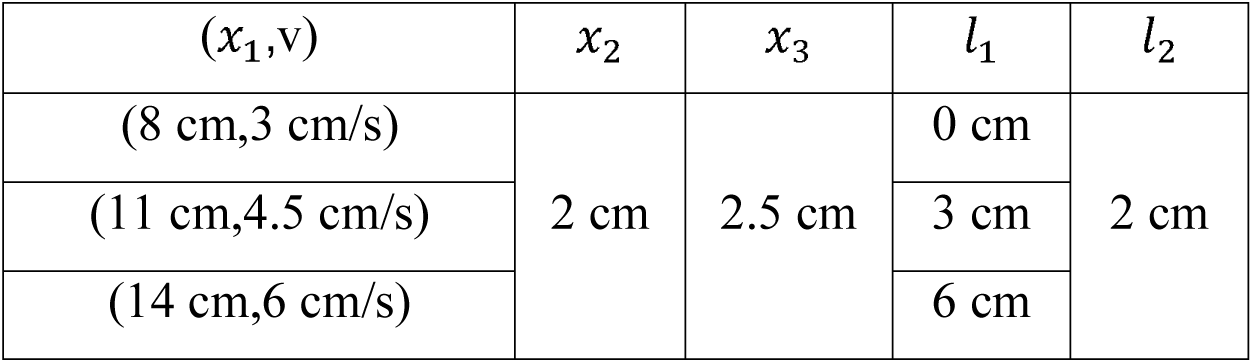
the parameters of the lateral paradigm.

**Figure 2.**
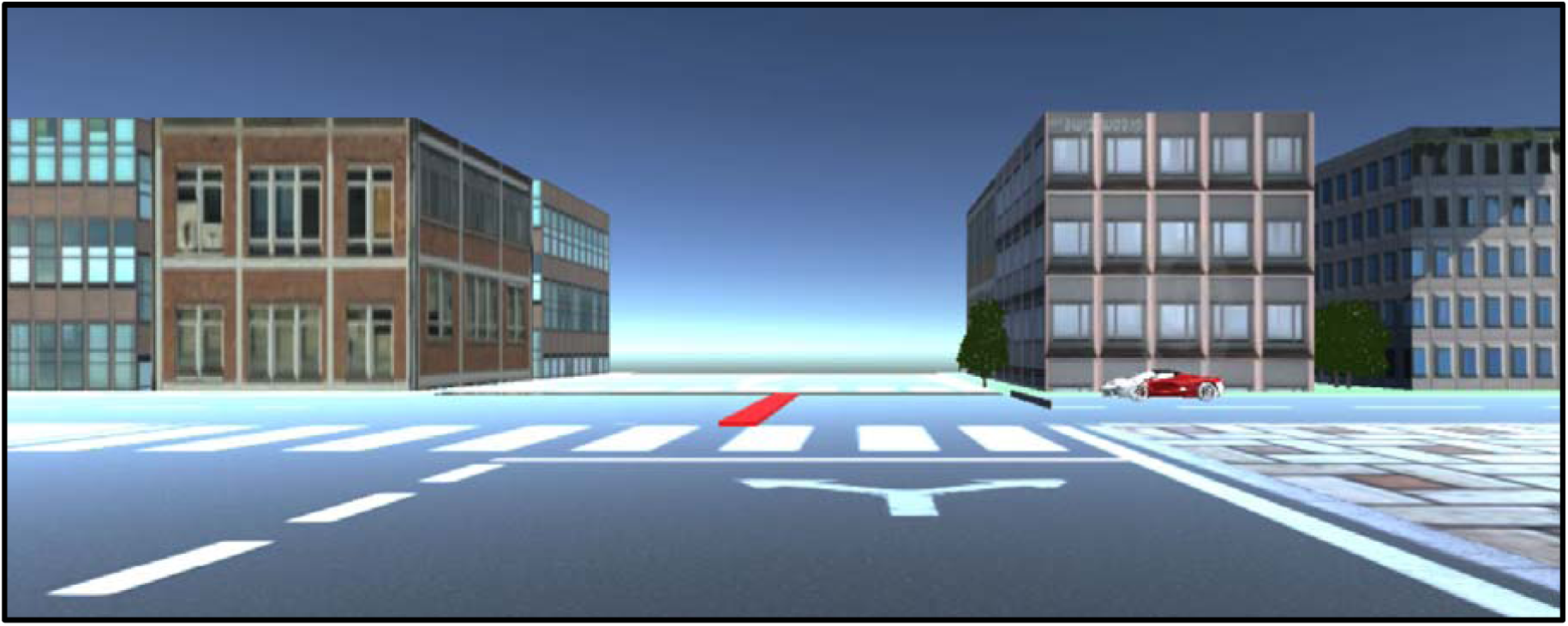
a screenshot of one of the trials in the lateral paradigm.

**Figure 3.**
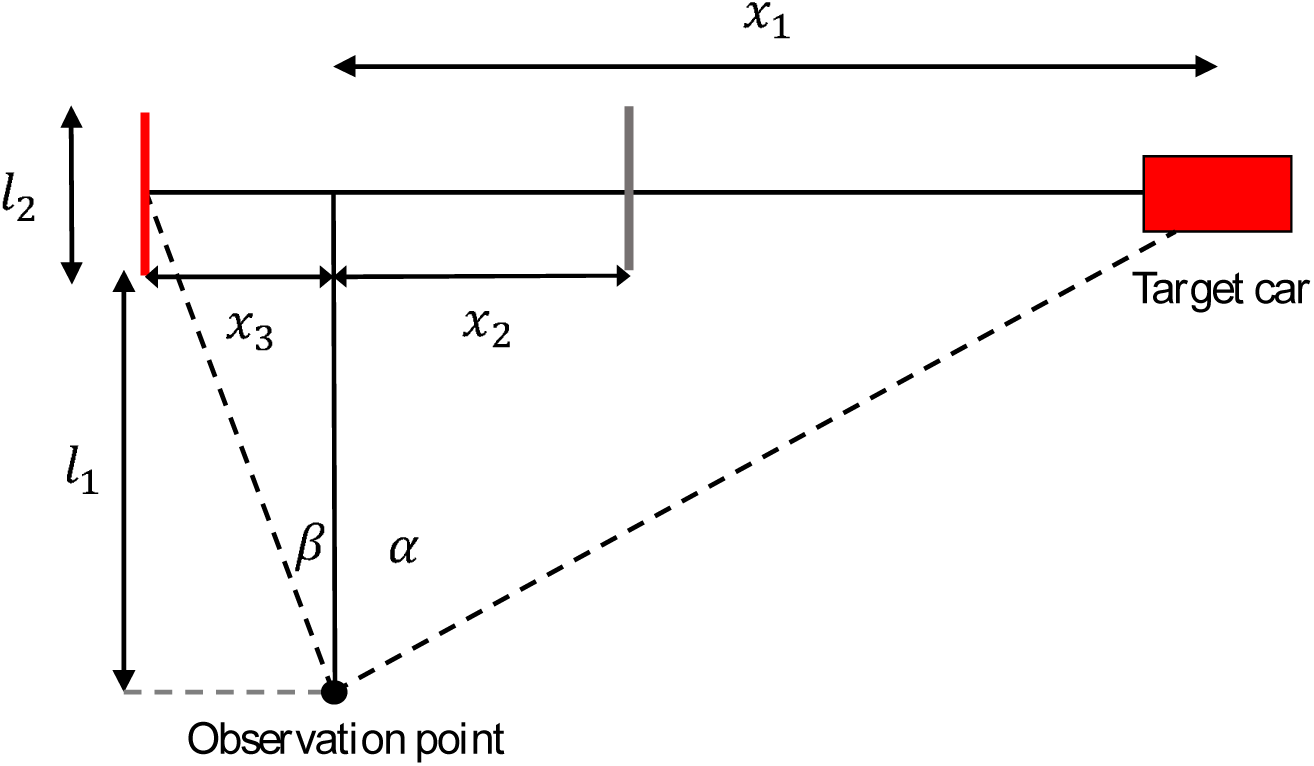
the schematic of lateral paradigm.

### 2.3. Experiment 2

After completing the lateral paradigm and taking a short break (ten minutes), participants were tested in a second experiment (hereafter called “approach paradigm”), in which all parameters were selected like the lateral paradigm, except that the target car moved towards the observation point instead of moving horizontally in frontoparallel plane.

**Figure 4.**
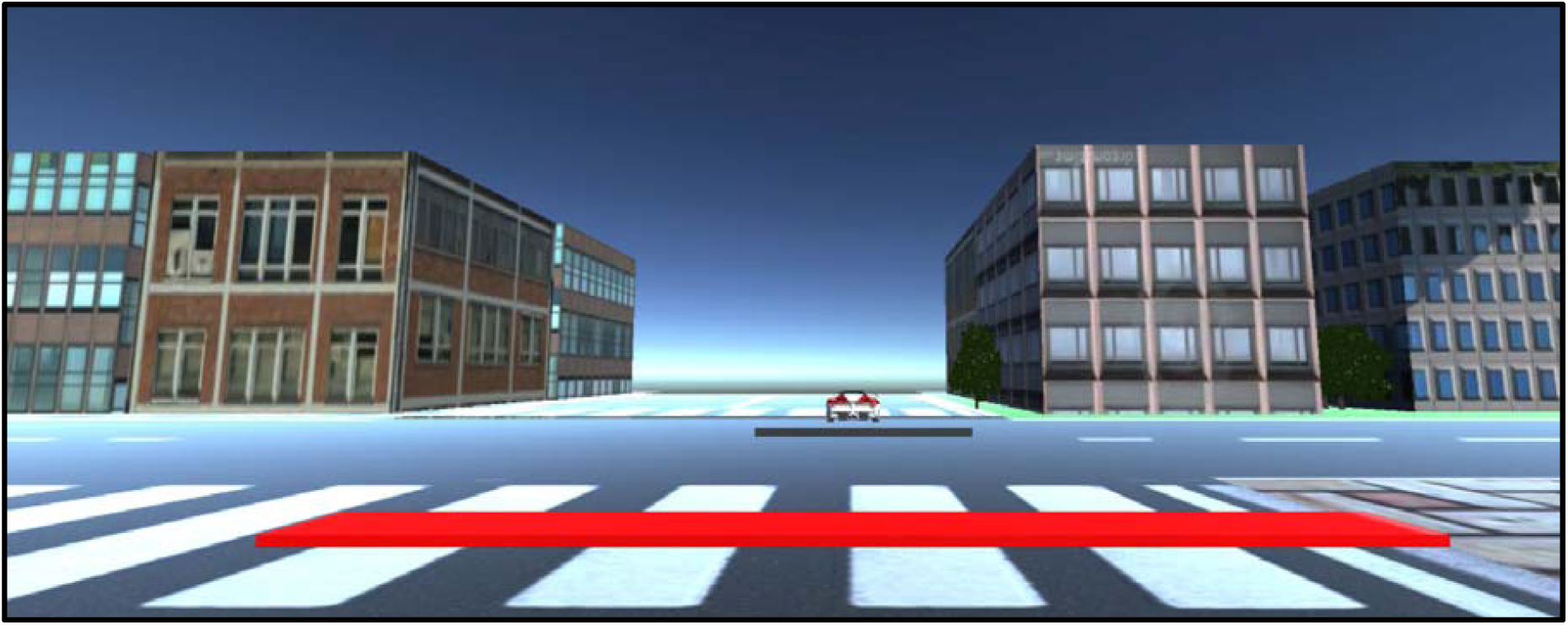
a screenshot of one of the trials in the approach paradigm.

Again, participants were asked to press a response key (down arrow key) at the instant when the car would have reached the finish line and no feedback on TTC estimation error was given to them. Figure 5 shows a schematic of the lateral paradigm. Table 2 shows the values of the parameters in the figure. There were nine combinations generated from three velocities for the target car and three distances between observation point and red line. Each trial was presented 10 times in random order, for a total of 90 trials.

**Table 2.** the parameters of the approach paradigm.

| $l_1$ | $(l_2, v)$ | $l_3$ | $x_1$ |
| --- | --- | --- | --- |
| 0 cm | (6 cm, 3 cm/s) | 4.5 cm | 2 cm |
| 3 cm | (9 cm, 4.5 cm/s) |  |  |
| 6 cm | (12 cm, 6 cm/s) |  |  |

**Figure 5.**
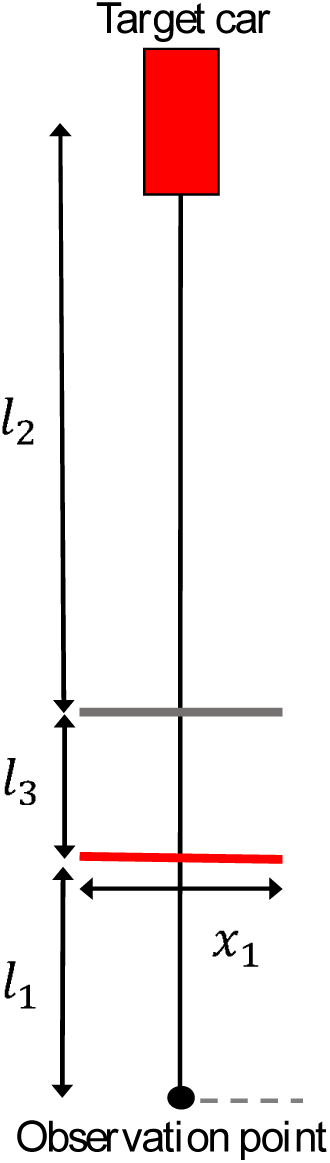
the schematic of approach paradigm.

## 3 Results and discussion

Performance of TTC-estimation in the present study was quantified through calculating error, defined by:

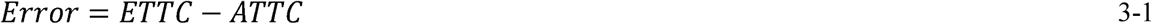

where *ETTC* (estimated TTC) is the time between the disappearance of the target car and the observer’s response, and *ATTC* (actual TTC) is the actual time between the disappearance of the target car and when it would have reached the finish line.

Consistent with the pattern of results reported in previous studies, judgment accuracy decreases with larger TTCs (Figure 6). In other words, decreasing the actual TTC results in decreasing the error in TTC estimation.

**Figure 6.**
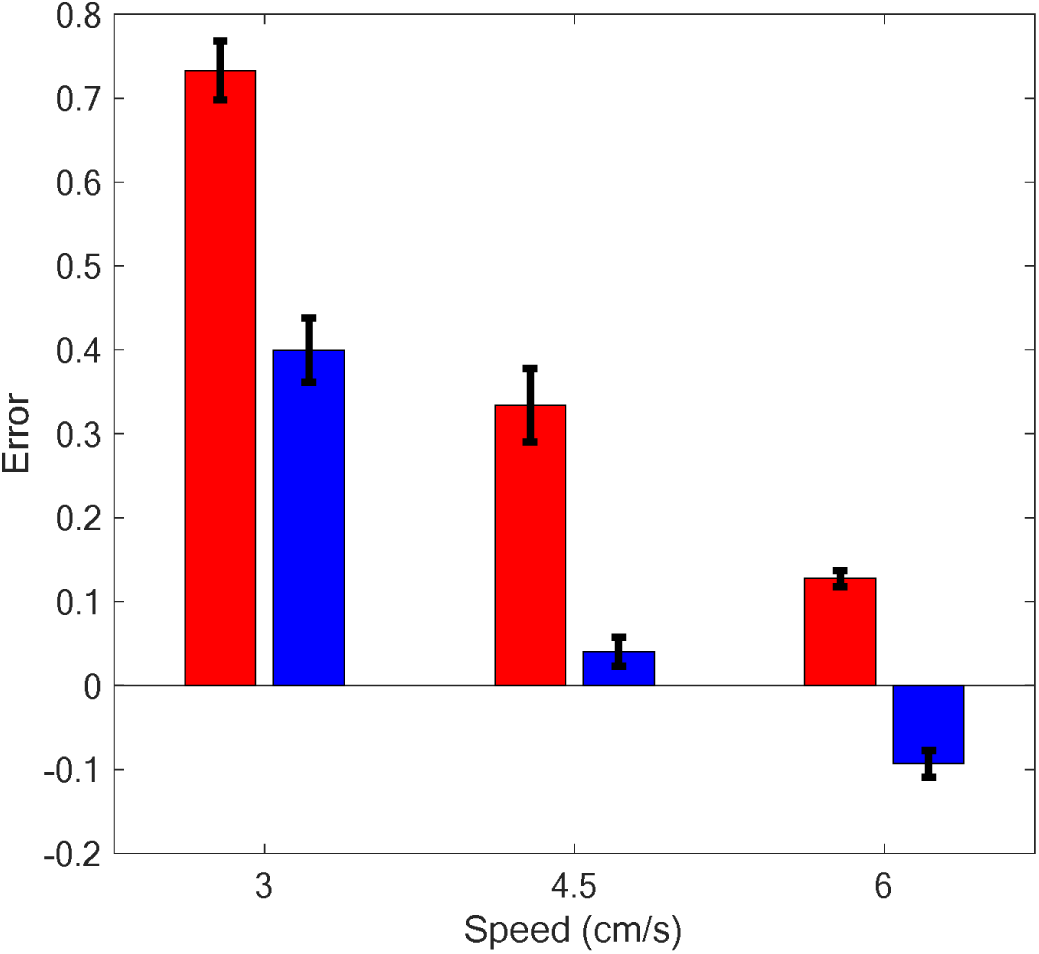
error values for the three levels of target car velocity, for lateral motion (Blue diagram) and for approach motion (red line). Horizontal axis shows three different velocities of 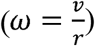 Vertical axis shows the error values.

In addition, estimated TTC decreases as actual TTC decreases (Figure 7). Furthermore, it is obvious for the approach motion the response times in three conditions are close to each other, meaning that participants were not able to understand the changes in the velocity of the target car properly.

**Figure 7.**
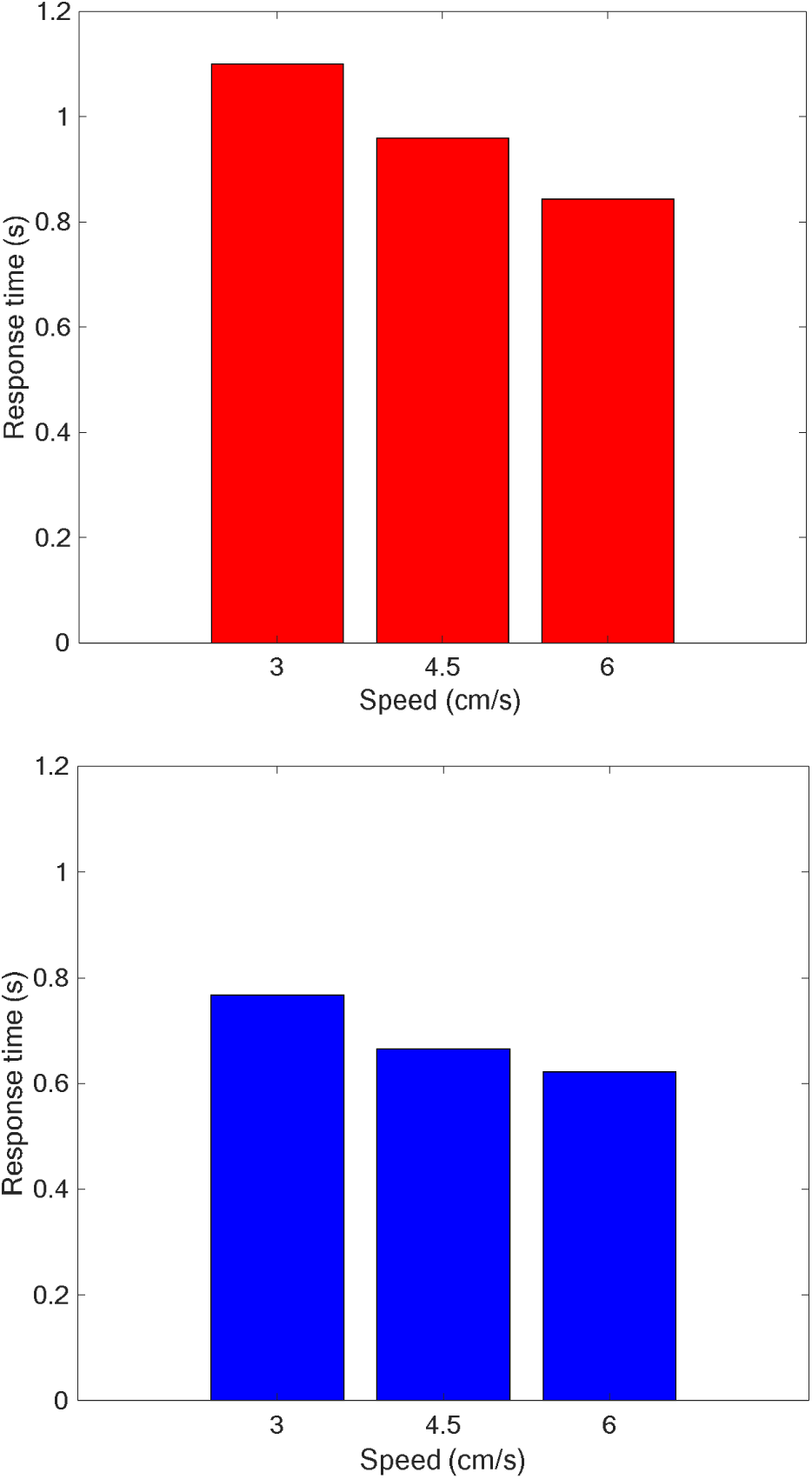
left: response times for the three levels of target car velocity, for approach motion; right: response times for the three levels of target car velocity, for lateral motion.

A three-way ANOVA was conducted on the *Error* values of all participants to examine the effect of the paradigm type, the velocity of the target car and the distance between observation point and contact point on TTC estimation. Paradigm types included two groups (lateral motion and approach motion), the velocity of the target car consisted of three levels (3 cm/s, 4.5 cm/s, 6 cm/s), and the distance between observation point and the intersection included three levels (0 cm, 3 cm, 6 cm). In all cases, the differences were statistically significant at the 0.05 significance level except for the distance between observation point and the intersection, meaning that the distance between the observer and the finish line did not make a difference in judging TTC in our experiment. The main difference obtained for paradigm type (Factor 1) yielded an F ratio of *F*(1,342)=174.812, *p*<0.001, indicating that the error is significantly higher for approach paradigm (*Mean*=0.037, *SD*=0.514) than for lateral paradigm (*Mean*=-0.419, *SD*=0.494). The difference obtained for the distance between observation point and intersection (Factor 2) yielded an F ratio of *F*(2,342)=0.943, *p*=0.391, indicating that the effect of this parameter was not statistically significant. Also, the differences obtained for the velocity of the target car (Factor 3) were statistically significant (*F*(2,342)=662.957, *p*<0.001; *Mean*_velocity1_=-0.95, *SD*_velocity1_=0.482; *Mean*_velocity2_=-0.919, *SD*_velocity2_=0.546; *Mean*_velocity3_=-1.020, *SD*_velocity3_=0.593). The pairwise interaction between parameters was not significant (the interaction between Factor 1 and Factor 2 yielded *F*(2,342)=1.393, *p*=0.250, the interaction between Factor 1 and Factor 3 yielded *F*(2,342)=0.391, *p*=0.677 and finally the interaction between factor 2 and factor 3 yielded *F*(4,342)=1.860, *p*=0.117).

According to motion parallax theorem, closer objects appear to move faster rather than objects that are farther away [20]. Therefore, it is expected that the distance between the observer and the intersection influence TTC judgment. Now, as shown in Figure 8, consider two objects moving in parallel trajectories from right to left (gray circles), while an observer (white circle) tracks their movements. For equal angular velocities, the farther object must move at a higher velocity than the nearer object (Figure 8-left), because for the same time-interval, the distance that the farther object should pass is larger than the distance that nearer object should pass.

**Figure 8.**
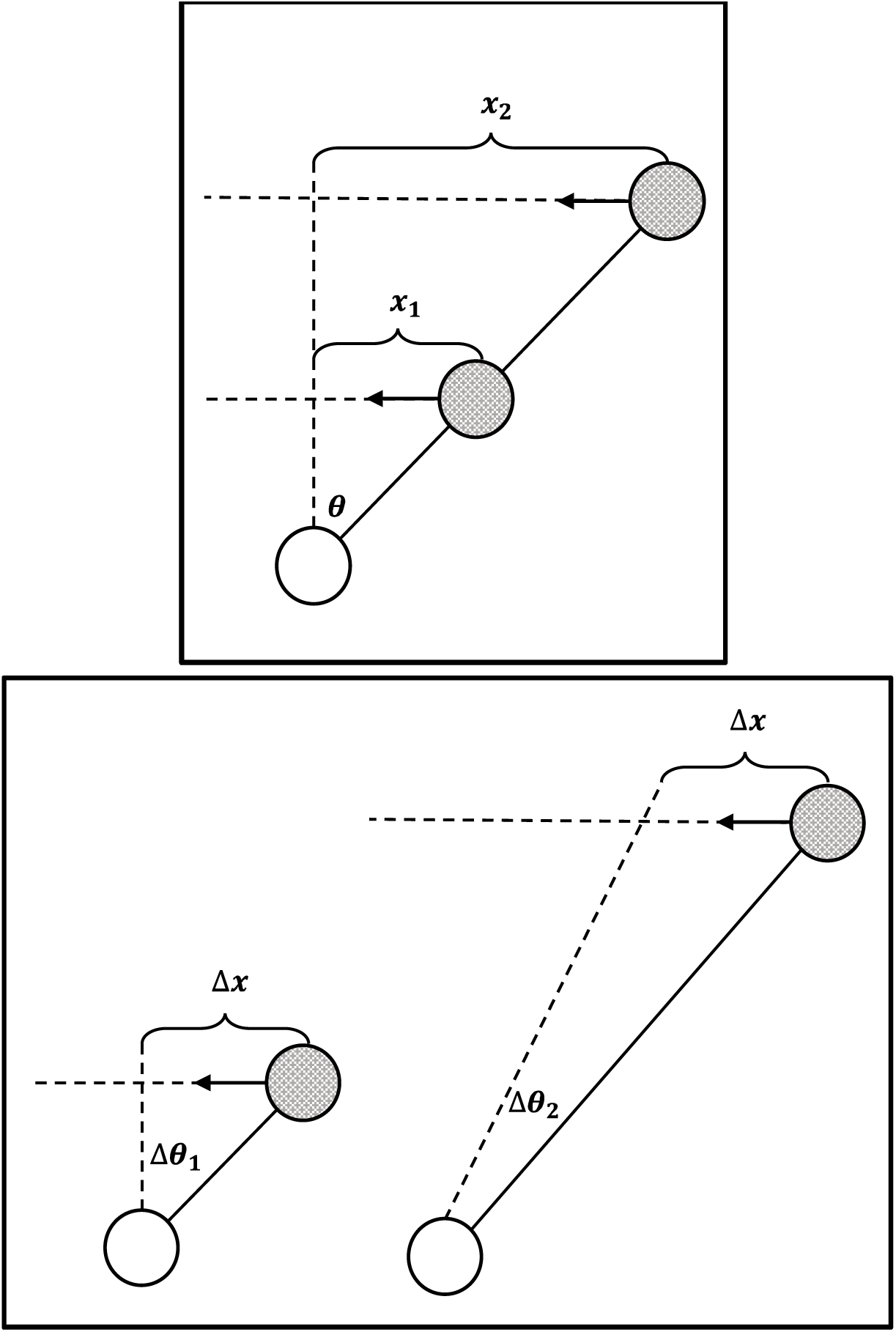
left: response times for the three levels of target car velocity, for lateral motion; right: response times for the three levels of target car velocity, for approach motion.

If these objects move at the same velocity, then angular velocity for the nearer object is more than the other object. For the same time interval and the same velocity, the angular velocity 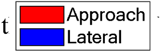 of the nearer object is larger than the angular velocity of the farther object.

where *v* is the velocity, ω is the angular velocity, and r is the distance between the observer and the moving object. It is obvious that for the farther objects r is larger, and at the same *v*, ω is smaller for the farther object. That is why closer objects appear to move faster than objects that are farther.

But, one may ask why changing the distance between the observer and the finish line did not cause a statistically significant difference in judging TTC in our experiment? According to Table 1

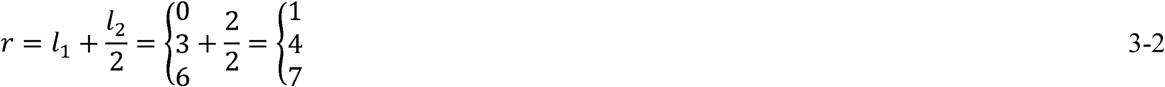

These values do not make significant difference between the angular velocities; therefore, motion parallax theorem cannot be understood through this experiment.

Now consider an object moving laterally from right to left at a constant velocity as shown in Figure 9. Points 1 through 5 represent five positions in its trajectory, which are marked at the same distance from each other. The red dot shows the observation point. Tangents of view angles can be written as:

**Figure 9.**
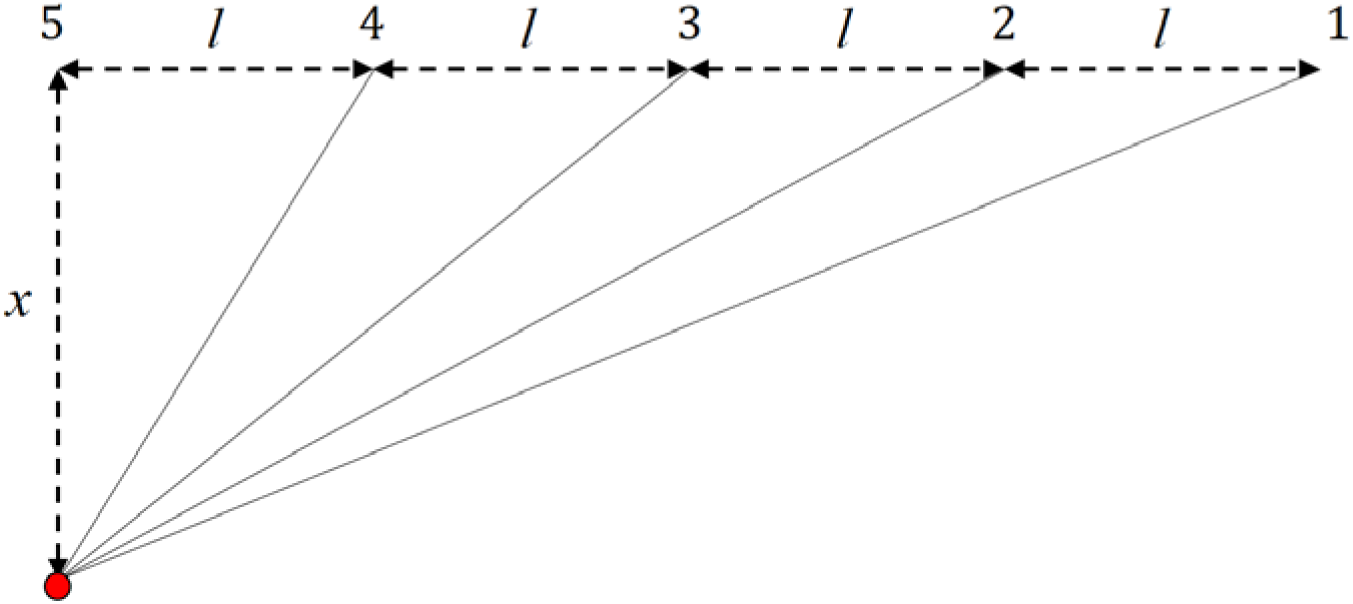
a schematic of five points of lateral motion which are positioned at the same distance from each other.

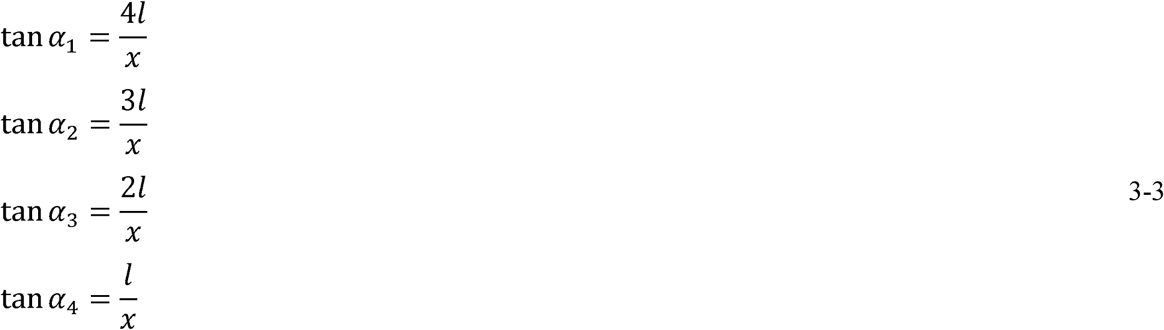

where α_1_, α_2_, α_3_, and α_4_ are visual angles for the points 1 to 4. When the object moves from one of these points to the next, the tangent of its visual angle changes according to the following equations:

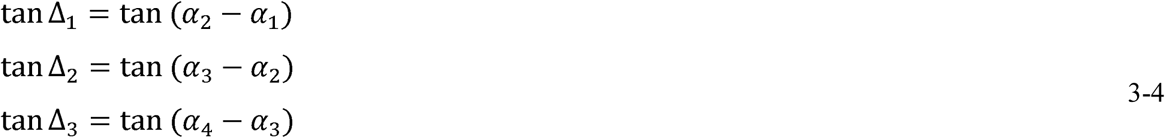

If the distance between two consecutive points is sufficiently small (*l* very small compared to *x*), then

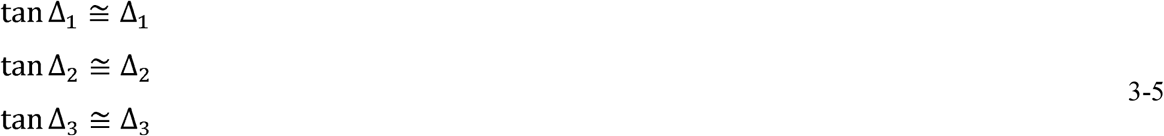

Therefore

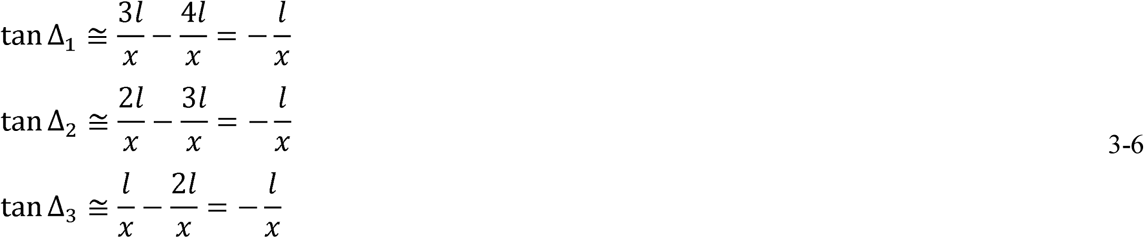

negativity indicates that the visual angle decreases as the objects moves from Point 1 to Point 5. Finally,

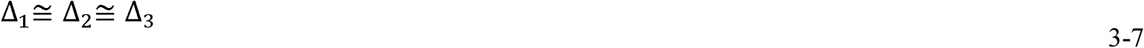

Meaning that visual angle changes at a constant rate, when the object moves laterally at a constant velocity.

Now, consider an object moving towards the viewer. Again five positions of its trajectory, having the same distance from each other are marked (points 1 to 5) and the red dot shows the observation point. Here, tangents of visual angles can be written as:

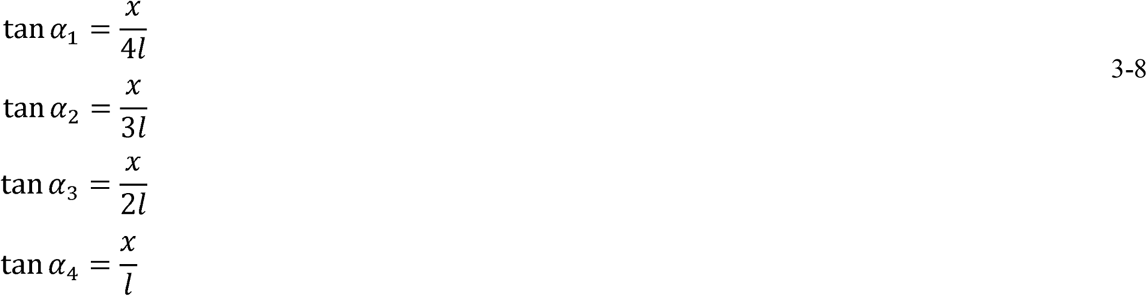

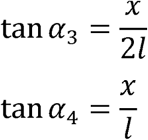

where α_1_, α_2_, α_3_, and α_4_ are visual angles for the points 1 to 4 and *l*s and *x* are remarked in Figure 10. When the object moves from one of these points to the next, the tangent of its visual angle changes according to the following equations:

**Figure 10.**
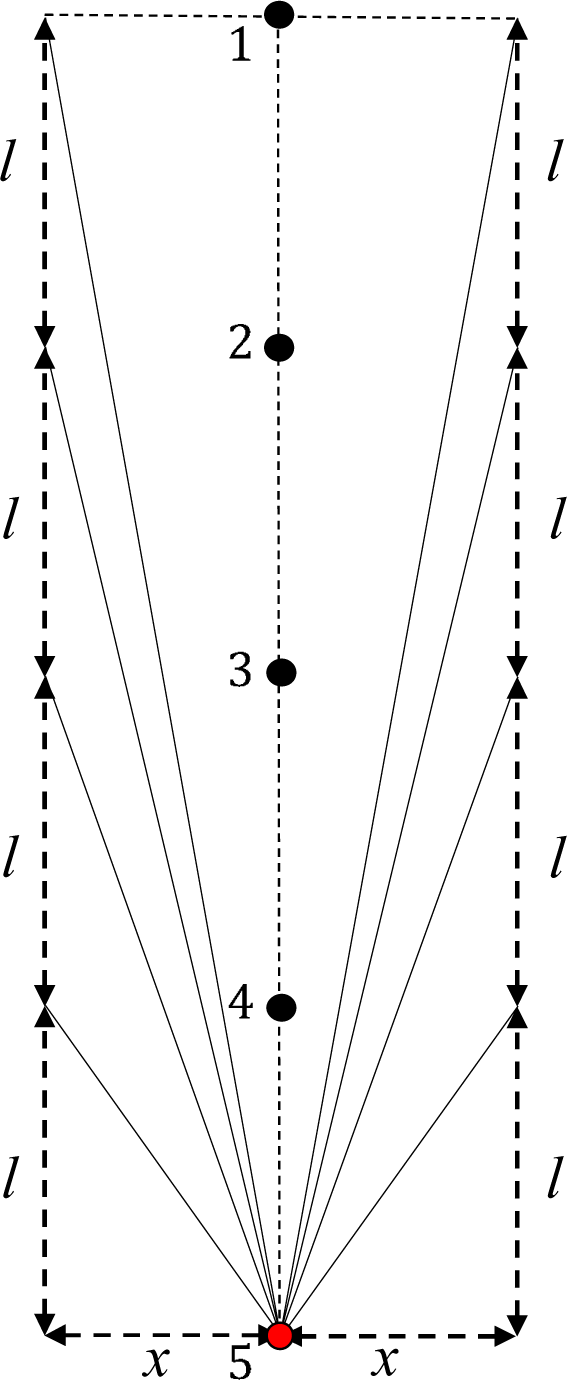
a schematic of approach motion. Points 1 to 5 are positioned at the same distance from each other.

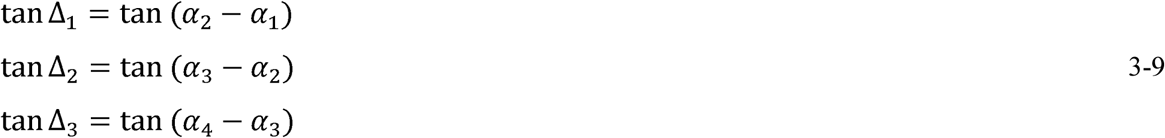

If the visual angles be small enough (*x* be very small compared to *l*), then

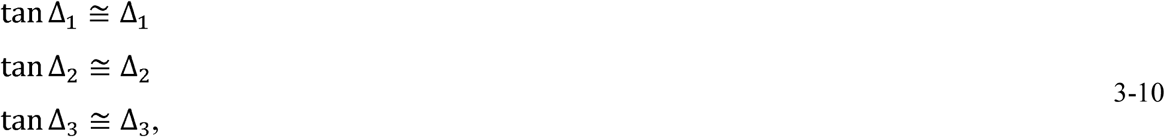

and

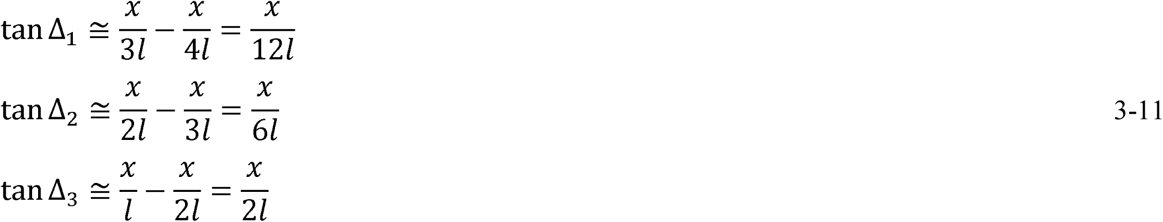

It is obvious that the visual angle increases as the object moves towards the observer. Therefore,

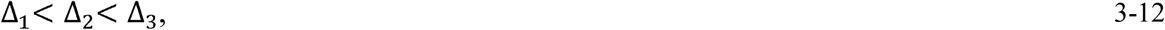

meaning that the rate of change in view angle is not constant when the object approaches the observer or moves away from it. As the object moves towards the observer, the angular velocity increases and the observer perceives an accelerated movement, while in the lateral motion, observer perceives a constant angular velocity. Humans are not able to estimate the acceleration value in moving visual stimuli [21, 22]. As a result, the performance for lateral motion TTC estimation is better than that of approach motion.

